# Stability Comparisons between Natural versus Engineered Archaeal Heat-Shock Proteins

**DOI:** 10.1101/2021.08.21.457208

**Authors:** Mercede Furr, Sreenivasulu Basha, Shilpi Agrawal, Zeina Alraawi, Piyasi Ghosh, Carson Stacy, T.K.S. Kumar, Ruben Michael Ceballos

## Abstract

Crenarchaeal group II chaperonins (a.k.a., “heat shock” proteins, HSPs) are abundantly expressed in species of the family *Sulfolobaceae*. HSPα and HSPβ expression is upregulated during thermal shock. HSPs are subunits of larger octadecameric complexes that function to protect intracellular proteins during thermal stress. Engineered HSPs have been constructed with the idea of protecting enzymes in industrial reactions. HSPβ-coh, a fusion protein comprised of HSPβ and type 1 cohesin from *Clostridium thermocellum* was used for *proof-of-concept*. Dockerin-endowed cellulolytic enzymes bind to the complex via cohesin-dockerin interactions. Enzymatic activity (i.e., hydrolysis of lignocellulose) is retained when the platform is used at high temperatures (e.g., 85-88°C). Moreover, enhancement persists on acid-pretreated substrates prompting the question: *Are HSPs acid tolerant?* In this study, HSP structural integrity is examined at different temperatures and pH. Far-UV circular dichroism and intrinsic fluorescence indicate HSPα and HSPβ retain structural integrity at neutral pH over a range of temperatures (25-90°C) while HSPβ-coh is less tolerant to thermal stress. Structural integrity is compromised for all subunits at ultra-low pH (i.e., pH 2) with HSPα showing the most susceptibility. Secondary structures of all HSPs are resilient under mildly acidic conditions (pH 4). ANS binding assays indicate a shift in tertiary structure for all subunits at ultra-low pH. Limited trypsin digestion reveals that the backbone of HSPβ-coh is the most flexible and HSPβ is the most resistant. Results suggest that HSPα and HSPβ are more resilient than HSPβ-coh under thermal challenge and that there are limits to the acid tolerance of all HSPs.

## Introduction

Chaperonin proteins form complexes that function to prevent misfolding and aggregation of other cellular proteins (1–3). Chaperonin complexes are typically large molecular cages (~1 MDa) that are upregulated during cellular stress (e.g., heat-shock). These molecular cages form ring structures (composed of one or more chaperonin subunit subtypes) that capture misfolded *client* proteins in an inner cavity (4). Refolding of misfolded substrates is accomplished by closing the chaperonin complex around the client protein. This chaperonin-client protein association perturbs local energy minima in the client protein in a process that involves hydrolysis of adenosine triphosphate (ATP) and divalent cation interactions (5,6). There are two groups of chaperonins. Group I chaperonins (e.g., GroEL/ES) are found in bacteria. The mechanism by which Group I chaperonins and their client proteins interact has been extensively studied (7–9). Group I chaperonins form tetradecamers by end-to-end stacking of heptameric rings thus forming a barrel-like structure within which protein may reside (10,11). GroEL requires a heptameric co-chaperonin, called GroES, to seal the barrel complex and assist in the refolding of the client protein (3,12). Group II chaperonins are found in archaea and eukaryotes. This includes the T-complex protein-1 ring complex (TRiC) or chaperonin-containing TCP-1 (CCT), which is found in eukaryotes. TriC/CCT consists of eight homologous subunits of distinct subtypes: CCT1-8 (13). Group II chaperonins expressed in archaea also form ring structures from subunits known as thermal factors (e.g., TF55, TF56); however, mechanisms of chaperonin-client protein interactions remain elusive. It is known that the archaeal heat shock protein complexes do not require a capping co-chaperone like the GroEL/ES bacterial system (14–16). It is also known that archaeal chaperonin complexes can include at least three chaperonin subtypes: HSPα (TF56), HSPβ (TF55), and HSPγ (17–20) with stoichiometry favoring HSPα/HSPβ complexes (4,21). However, the most distinguishing feature of these archaeal chaperonins is that those found in hyperthermophilic archaea, such as those of the Order Sulfolobales, function at temperatures exceeding 74°C. Indeed, multiple species within the family *Sulfolobaceae* have been isolated from sulfuric geothermal pools and volcanic hot springs at temperatures from 74°C-88°C and pH 1-4 (17,19). Although some group II chaperonin complexes can form heptameric or octameric ring structures, chaperonin complexes from Sulfolobales tend to form double nonameric-ring structures (17,18,22–25). It has also been shown that complex formation and subunit expression are upregulated in response to thermal shock (18,20). In *in vivo* and *in vitro* experiments HSPα/HSPβ admixtures appear to dominate these heat-shock protein complexes (17,18,24). It has been suggested that HSPγ may be a “cold-shock” protein since its incorporation into chaperonin complexes *in vitro* is actually favored at lower temperatures in the physiological range (14). Studies have shown that HSPα homomers, HSPβ homomers, and HSPα/HSPβ heteromers will form *in vitro* in the presence of ATP and Mg^2+^. Yet, functional implications of homomeric versus heteromeric complexes and the significance of alternative HSPα:HSPβ stoichiometries *in vivo* remain unclear. Whether HSP complexes are homomeric or heteromeric during thermal shock, subunits appear to work together in a highly cooperative manner (1,3,26). It is suggested that function and substrate specificity of archaeal chaperonins is mediated by subunit composition and complex geometry (27).Thus, characterizing the biophysical properties of HSP subunits which form chaperonin complexes may elucidate key structure-function relationships that mediate HSP complex-client protein dynamics under varying environmental conditions (e.g., thermal shock or pH flux).

Due to the thermotolerance of crenarchaeal group II chaperonins, efforts to use these heat shock proteins in biotechnology has resulted in the design of engineered platforms including complexes composed of subunits which are fusion constructs between HSPs and accessory proteins (28–31). One such effort features the fusion of cohesin (Type I) from the bacterium *Clostridium thermocellum* to a circular permutant of HSPβ derived from *Sulfolobus shibatae* (28,30). The resulting HSPβ-cohesin (HSPβ-coh) fusion protein was shown *in vitro* to form double nonameric-ring (18-mer) complexes resembling natural thermosomes. This engineered prototype complex was called a *rosettasome* (28) and was used to bind cellulases from *C. thermocellulum* via cohesion (fused to HSPβ) and dockerin (Type 1) found as a functional domain on cellulolytic enzymes of the *C. thermocellum* cellulosome. The rosettasome bound with cellulolytic enzymes (a.k.a., the rosettazyme) was shown to more efficiently breakdown Avicel® than free enzyme in solution absent the chaperonin-based platform (28).

Thereafter, this engineered construct was shown to improve hydrolytic efficiency on acid- and alkaline-treated lignocellulosic biomass (30). Although catalytic activity of enzymes bound to the engineered platform resulted in a 2- to 3-fold increase in hydrolysis of carbohydrates (i.e., cellulose, hemicellulose) on pretreated corn pericarp and wheat straw substrates (30), a technology transfer assessment concluded that a 6- to 10-fold enhancement of enzymatic efficiency is required for economic viability. This level of enhancement could not be accomplished solely by varying enzyme complement. Therefore, the core platform (i.e., engineered complex) required re-design, testing, and optimization. This resulted in the construction of a next-generation *mobile enzyme sequestration platform* (MESP)

As in prior work, MESPs appear to enhance enzymatic efficiency on acid-pretreated substrates. Although the thermotolerance of engineered platforms was anticipated, the enhanced enzymatic efficiency on acid-treated substrate was not. This result suggested that MESPs may be acid-tolerant. Despite surviving in acidic geothermal habitats at pH<4, the intracellular environment of Sulfolobales is reported to be at a pH of ~6.5 (32,33). Therefore, there was no expectation of HSP acid tolerance. In the present study, we analyze the relative stability of natural HSPs (HSPα and HSPβ) and the engineered construct (i.e., HSPβ-coh) under different conditions of pH and temperature using a variety of biophysical methods to determine the limits of the acid and thermal tolerance of these proteins.

## Results

### Silver staining confirms purity of HSP protein working stocks

Using sodium dodecyl sulphate-polyacrylamide gel electrophoresis (SDS-PAGE) and silver staining, purified HSPα, HSPβ, and HSPβ-coh subunits were compared to a broad-range pre-stained ladder. HSPα and HSPβ produce ~60 kDa bands and the larger HSPβ-coh fusion construct produces a band at ~73 kDa (Fig. S1), which is consistent with the molecular weights derived from the ExPASy ProtParam server (34). HSPα and HSPβ differ, in terms of length, by 8 residues, with 560 and 552 amino acids, respectively. The molecular weight of the engineered HSPβ-coh fusion protein is higher at 73 kDa with 660 amino acids. Mass spectroscopy (i.e., MALDI-TOF MS) was employed (Fig. S2) to confirm the primary structure of all HSPs (i.e., amino acid sequences).

### Modeling predicts structural organization of HSP subunits

Upon resolving the primary structures of each of the HSP subunits (i.e., HSPα, HSPβ, and HSPβ-coh), the primary sequences were used to generate structural models (see Fig. 1) in PyMOL 2.4 (https://pymol.org) (35). Structural models for HSPα and HSPβ exhibit similar three-dimensional structures (Fig. 1A, B). These data were used to compare secondary structures (i.e., α-helices, β-sheets, loops, and turns), hydrophobic segments, conserved regions, and three functional domains (i.e., apical, intermediate, and equatorial domains). In general, all three HSP subunits appear to be helix-rich structures with 16 α-helices, 18 β-strands, and a characteristic stem loop between the intermediate and equatorial domains, which is consistent with higher resolution models previously reported (4). These domains show structural similarities to the GroEL superfamily of group I chaperonins in bacteria (27). Results from multiple sequence alignments of the engineered HSPβ-coh show 42% and 66% sequence similarity with the natural HSPα and HSPβ chaperonins, respectively. Out of the 560 amino acids that comprise natural HSPα, it is estimated that 52.68% are found in α-helices and 14.29 % in β-strands. Likewise, out of the 552 amino acids that comprise natural HSPβ, it is estimated that 53.08% are found in α-helices and 14.49% in β-strands. Apical domains for HSPα and HSPβ each resolve from a contiguous stretch of residues. However, the equatorial and intermediate domains for HSPα and HSPβ form from the convergence of multiple stretches of amino acids and the resulting intermingled secondary structural elements.

**Figure 1.**
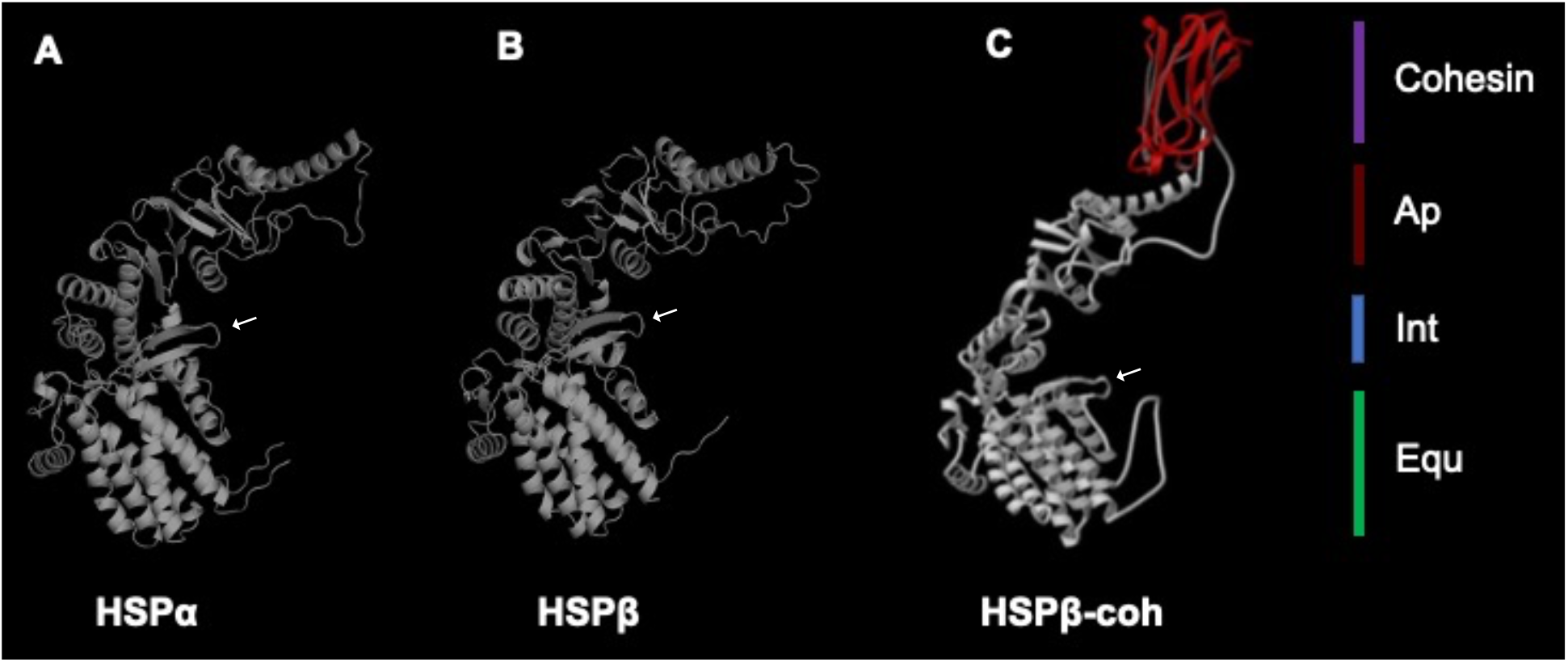
Structural models of natural and engineered HSP subunits. (A) 3-D structural model of the HSPα subunit; (B) structural model of HSPβ; (C) structural model of the engineered HSPβ-coh (adapted from Mitsuzawa et al., 2009). Homologous apical (maroon), intermediate (blue), and equatorial (green) domains are shown by section bars (right). Characteristic stem loop between equatorial and intermediate domain is present on all the three subunits (white arrow).

All HSP subunits exhibit comparable tertiary structures. The base structure of HSPβ-coh resembles HSPα and HSPβ but includes the cohesin extending from the apex of the HSPβ permutant (Fig. 1C). Each model reveals a three-dimensional structure with apical, intermediate, and equatorial domains. The cohesin domain fused to the HSPβ circular permutant (i.e., HSPβ-coh) provides an interaction site for dockerin (type 1) found on *C. thermocellum* cellulases, xylanases, and dockerin-endowed engineered fusion protein (i.e., enzyme) constructs (36). Since HSPβ-coh is based on the fusion of cohesin to a circular permutant of HSPβ, the location of residues comprising key secondary structure elements is shifted. Still, the number of α-helices and β-strands (as well as their relative positions in the tertiary structure of the folded protein) remains conserved. HSPβ-coh exhibits 16 α-helices, 18 β-strands, and the aforementioned characteristic HSP stem loop. In HSPα, the stem loop resolves at residues 46-58 (KMLIDSFGDVTIT). In HSPβ, the stem loop is located from positions 56-68 (KMFVDSLGDITIT). In HSPβ-coh the stem loop resolves at residues 315-327(KILVDSLGITIT). This HSP stem loop structure, which is also present in group I chaperonins (i.e., GroEL/ES), features an ATP binding site, which is critical for chaperonin function (37).

Alignment of HSPα, HSPβ, and HSPβ-coh reveals other highly conserved stretches of amino acid sequences between 57 - 73, 36-48 and 85-93 for HSPα, 95-103, 46-58 and 67-83 for HSPβ and 355-362, 316-332 and 325-341 for HSPβ-coh (Fig. 2), including notable hydrophobic regions. As part of the HSPβ-coh fusion construct two linkers are present in the primary structure (Fig. 2, orange icons). One linker at position 265-270 was engineered to alter the position of the N-terminus and C-terminus in the development of the permutant. The second linker shown at position 526-534 was added to fuse cohesin (type 1) from *C. thermocellum* to the *S. shibatae* HSPβ permutant. For experiments in this study the 9-residue linker form of HSPβ-coh was used. However, other linker lengths are also available (*data not shown*). Likewise, other HSPβ circular permutants are available. Changing the position of cohesin using different circular permutants and the HSPβ-coh linker length provides opportunities to study HSP complex-client protein interactions and the role of linker length in preventing steric interference between enzymes bound to the MESP. (These alterations will be reported separately).

**Figure 2.**
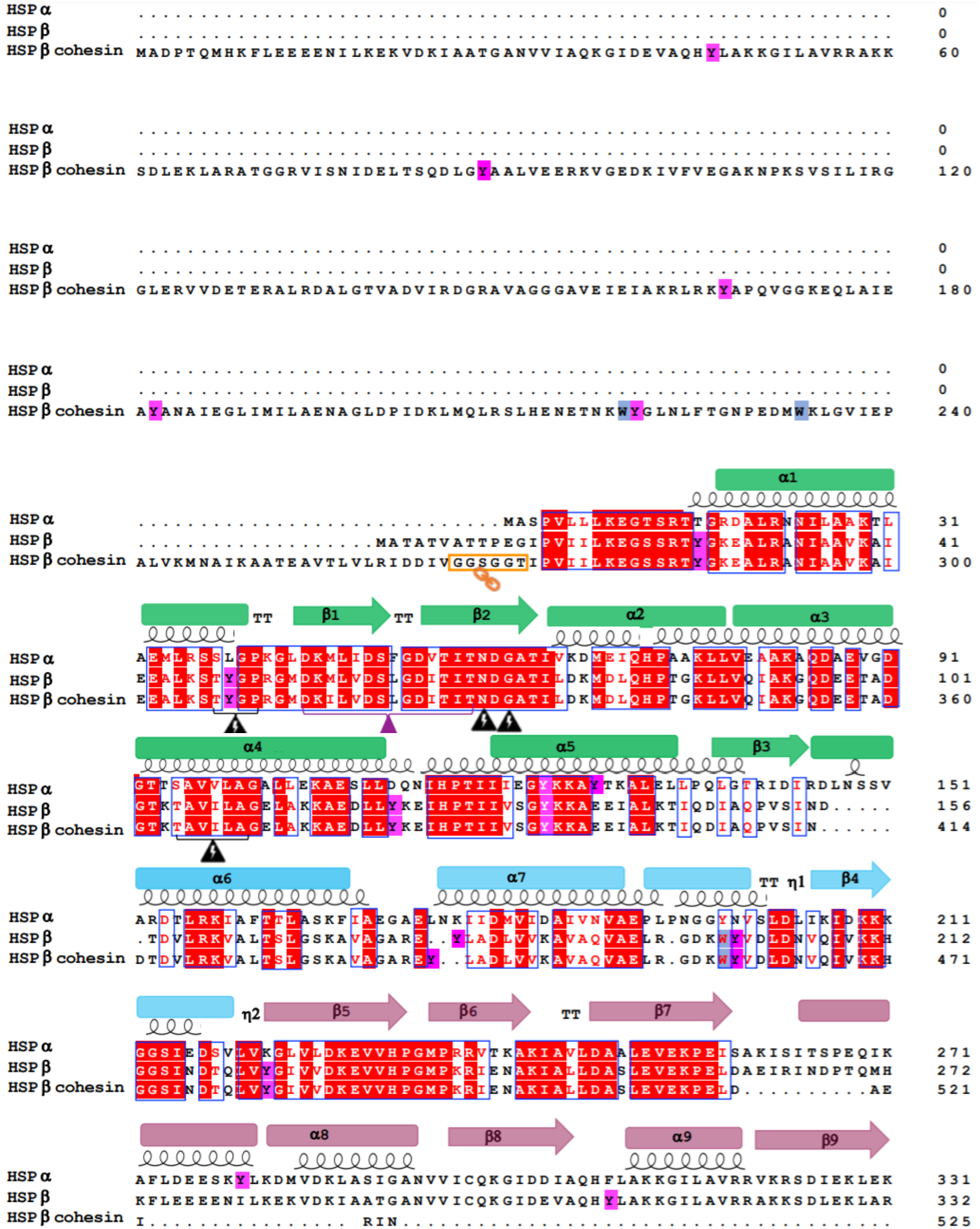
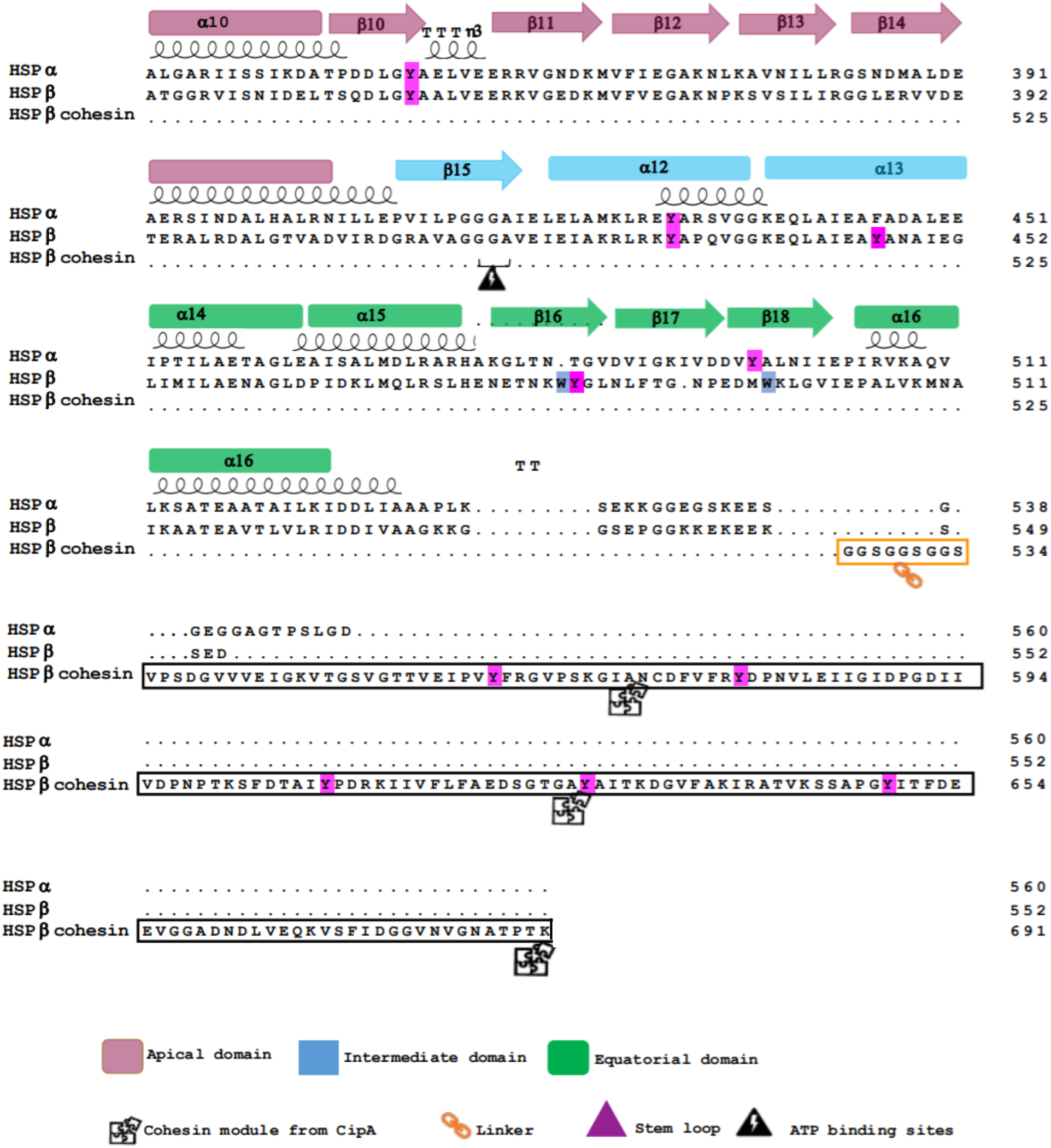
Primary sequence alignment of HSPα, HSPβ, and HSPβ-coh. Multiple sequence alignment was generated using ClustalW (38) and visualized by ESPript (39).The α-helices and β-strands are represented as coils and arrows, respectively. β-turns are marked noted as TT. Conserved regions are marked in blue boxes and the residues are highlighted in red. The linker at the amino acid position 267 of HSPβ and between HSPβ and the cohesion moiety is marked with an orange box. The domains are color coded: apical (magenta), intermediate (blue), and equatorial (green). The stem loop region is denoted by a purple triangle. Tyrosine residues are highlighted in purple and tryptophan residues are highlighted in blue.

### Differential Scanning Calorimetry (DSC) reveals high thermal stability of HSP subunits

DSC is an analytical tool commonly used for determining the thermal stability of proteins by measuring changes in heat capacity (40). DSC was employed to assess the limits of the HSP subunit thermal stabilities. Specifically, HSPα, HSPβ, and HSPβ-coh were each subjected to temperatures from 25°C −110°C. Readings were recorded at 5°C increments. The melting temperature (T_m_) at which 50% of the protein exists in denatured or unfolded states was calculated. T_m_ thresholds of HSPα, HSPβ, and HSPβ-coh were determined under different pH conditions (2, 4, and 7). T_m_ for HSPα, HSPβ, and HSPβ-coh at pH 7 were determined to be 93.6°C, 93.6°C, and 88.3°C, respectively (Fig. 3). On the contrary, T_m_ of HSPα at pH 2 and 4 shifts from 93°C to 68.2°C and 69.4°C, respectively, whereas T_m_ of HSPβ at pH 2 and 4 shifts from 93.0°C to 58.5°C and 57.0°C, respectively (Fig. 3). T_m_ of HSPβ-coh shifts from 88.3°C at pH 7 to 67.5°C and 66.0°C at pH 2 and 4, respectively (Fig. 3).

**Figure 3.**
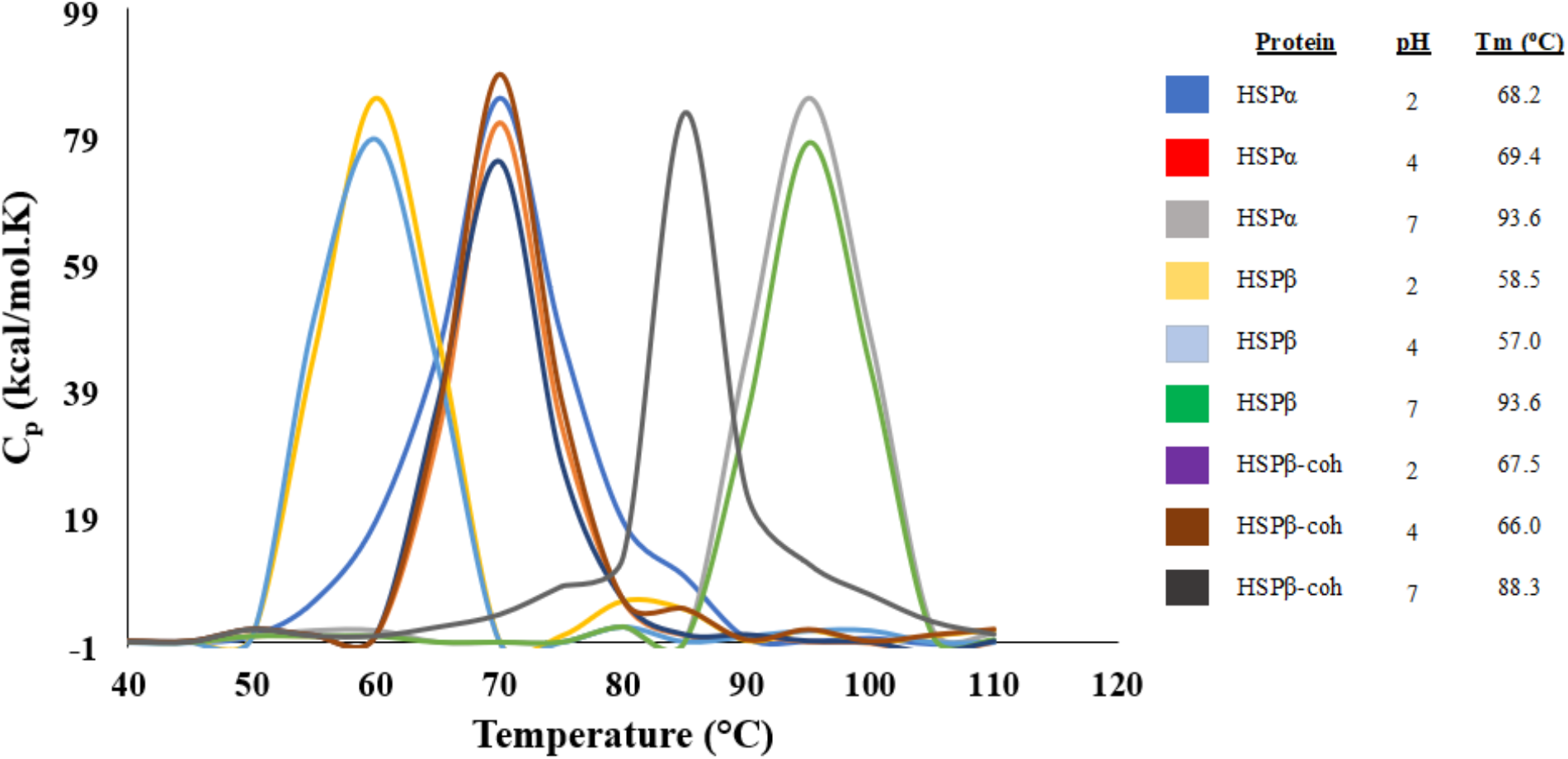
Differential Scanning Calorimetry of HSP subunits. Analysis of the thermodynamic stabilities of HSPα, HSPβ, and HSPβ-coh monitored by DSC under three different pH conditions: HSPα pH 2 (red), HSPα pH 4 (purple), HSPα pH 7 (orange), HSPβ pH 2 (magenta), HSPβ at pH 4 (green), HSPβ pH 7 (cyan), HSPβ-coh pH 2 (brown), HSPβ-coh pH 4 (blue), and HSPβ-coh pH 7 (black).

### Integrity of the secondary structures of HSP subunits at different temperatures

To determine the stability of secondary structural elements (e.g., α-helices) under varying conditions of pH and temperature, far-UV circular dichroism (CD) spectroscopy experiments were conducted for each subunit. The CD spectra for each HSP subunit reveals two hypo-elliptical bands around 208 nm and 222 nm (see Fig. 4), which is characteristic of helical-rich protein backbone structures (41,42). The changes in circular dichroism spectra have been examined as a function of temperature and pH. At neutral pH (pH 7), varying temperature does not significantly impact the secondary structure of HSPα, HSPβ or HSPβ-coh (Fig. 4A). The thermal denaturation CD spectral overlays of HSPα, HSPβ, and HSPβ-coh at pH 7 superimpose with one another with negligible deviation (Fig. 4A). Since HSPs have evolved in thermophilic environments with intracellular pH typically around 6.5-6.8 (32,33), it is suggested that neutral pH trials at the higher end of the temperature range (75°C-80°C) best represent the native structural state. The magnitude of molar ellipticity decreases as temperature increases at pH 2 and 4 for HSPα and at pH 2 for HSPβ-coh (Fig. 4B-C). Minor spectral shifts along the x-axis (i.e., wavelength) were also observed at pH 2 and 4 for HSPα (Fig. 4C).

**Figure 4.**
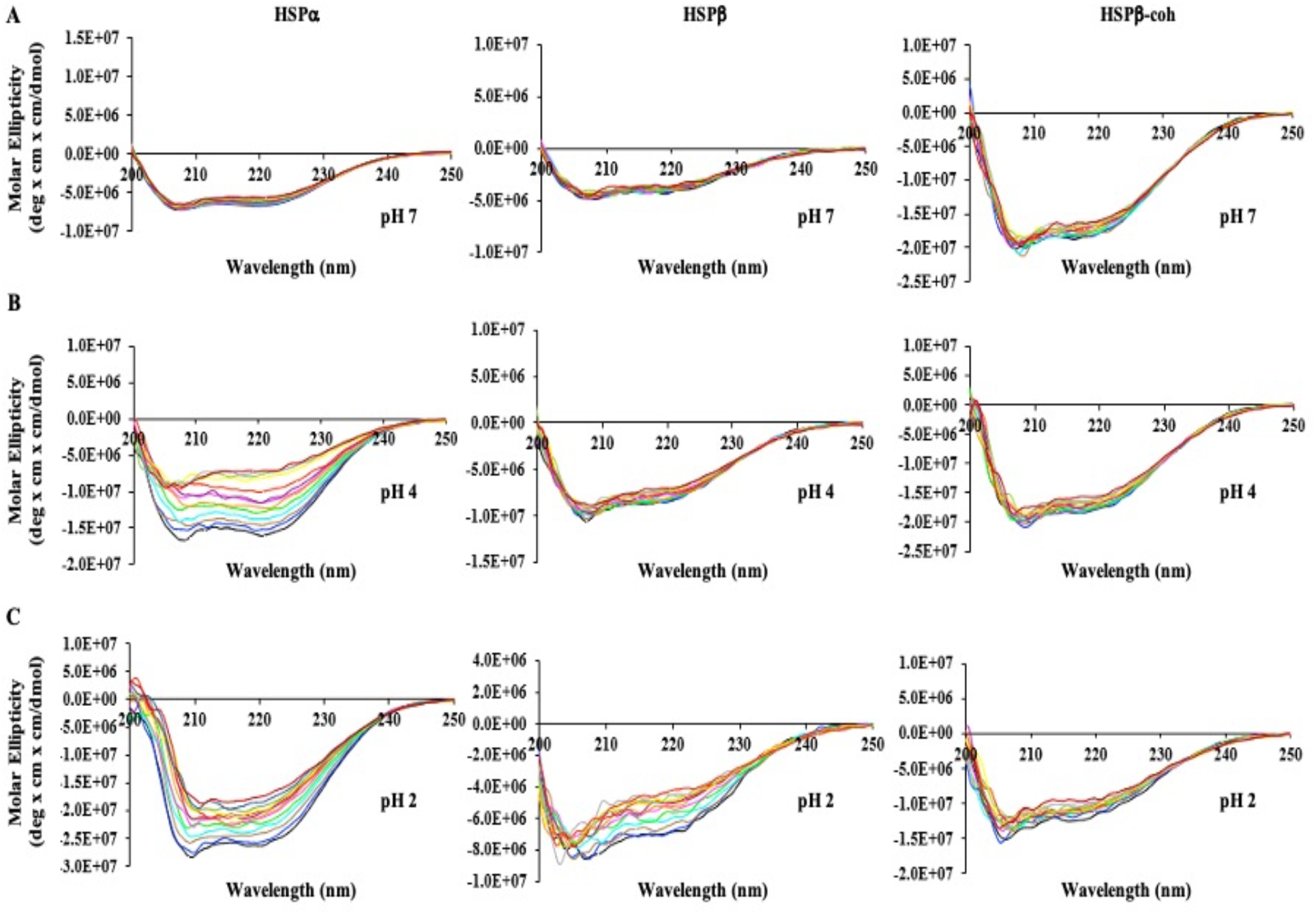
Far-UV Circular Dichroism (CD) spectra of HSPs. HSPα (Left), HSPβ (Middle), and HSPβ-coh (Right) upon thermal denaturation (25-90°C) at (A) pH 7, (B) pH 4, and (C) pH 2. Color coding is the same for each spectrum overlay: 25°C (black), 30°C (blue), 35°C (brown), 40°C (teal), 45°C (neon green), 50°C (pink), 55°C (orange), 60°C (purple), 65°C (red), 70°C (yellow), 75°C (green), 80°C (grey), 85°C (maroon).

Calculating the weighted spectral difference (WSD) provides a detailed method for quantifying spectral differences in CD data (43). WSD was employed to provide a more precise comparison of the spectral similarities and dissimilarities between subunits under varying conditions. Again, we consider the spectra at pH 7 and 75°C to be representative of the native conformation for each HSP subunit. All subunits show similarity across the temperature range of 25-90°C at pH 7. This is represented by the overlap of WSD values in blue (Fig.5). Interestingly, the thermal denaturation spectra of HSPβ-coh at pH 4 also overlaps with the representative native conformation spectra of all three subunits at pH 7. However, the thermal denaturation spectra of HSPβ at pH 4 and 2 reveal a decrease in magnitude. Although the WSD values derived from the thermal denaturation spectra of HSPα deviates the most from the native conformation of itself and the other subunits at pH 2, all the subunits, at pH 2, show dissimilarity when compared to the native conformation spectra. (Fig.5, indicated in red). Analysis of WSD across temperature and pH ranges indicate changes in pH have a larger effect on the secondary structure of each HSP than changes in temperature (Fig. 5). One exception to this trend was seen with HSPβ-coh at pH 4 closely matching the trend curves at pH 7 across the range of temperatures. A negative correlation was seen between temperature and WSD for HSPβ in low pH conditions. These results suggest that, in terms of secondary structural stability, HSPα is more sensitive to changes in pH and temperature than HSPβ or HSPβ-coh. Despite apparent sensitivity to low pH, results suggest thermal stability for HSPα, HSPβ, and HSPβ-coh up to 85°C.

**Figure 5.**
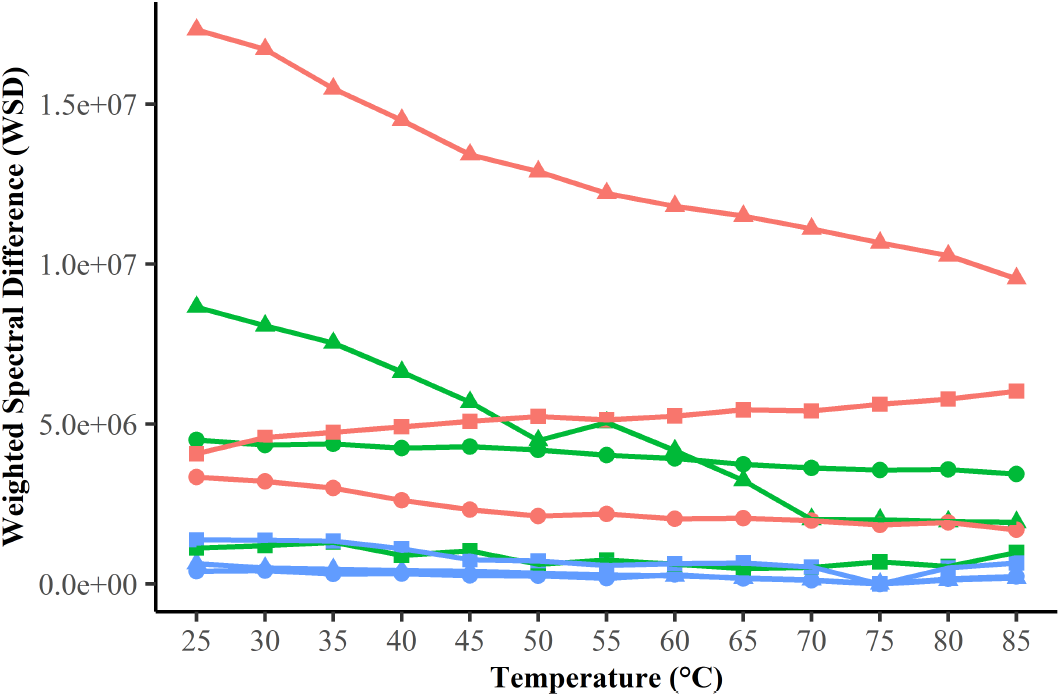
Weighted Spectral Difference (WSD). WSD of Far-UV Circular Dichroism (CD) spectra for HSPs. HSPα (▲), HSPβ (•), and HSPβ-coh (□) at pH 2 (red), pH 4 (green), and pH 7 (blue). Each point indicates the calculated WSD between the spectra of a given combination of protein, pH and temperature and the spectra of the same protein at pH = 7 and T = 75°C.

Although CD spectra for HSPα exhibited the most significant decrease in magnitude at lower pH, the general spectral profile remained. To determine if these shifts represent significant changes in secondary structure, the 222/208 nm ratio was plotted for each spectrum as a function of temperature (by pH condition) for each HSP (Fig. 6). The ratio of the molar ellipticities at 222 nm and 208 nm ([θ]222/[θ]208) is utilized as a criterion to evaluate the presence of coiled-coil helices in proteins (44). A non-interacting α-helix presents a ratio at 0.9 or below, while ratios of 1.0 or above indicate the presence of coiled-coil helices (45,46). The effect of varying the temperature and pH of the natural and engineered HSP subunits on the [θ]222/[θ]208 ratio was examined, to determine if coiled-coil formation occurs due to fluctuations in temperature and pH. The ratio remained unaltered (below 0.9) for all three subunits at pH 7 over temperatures ranging from 25-85°C (Fig. 6). There is no overt alteration in the ratio for HSPβ and HSPβ-coh in the lower pH range (2 and 4) (Fig. 6). For HSPα, induction of coiled-coil helices appears to take place over the temperature range of 40-85°C at pH 2 as indicated by a shift in 222/208 nm to 1.0 (and above). The emergence of coiled-coil structures within HSPα at pH 4 appears to take place from 30-65°C.

**Figure 6.**
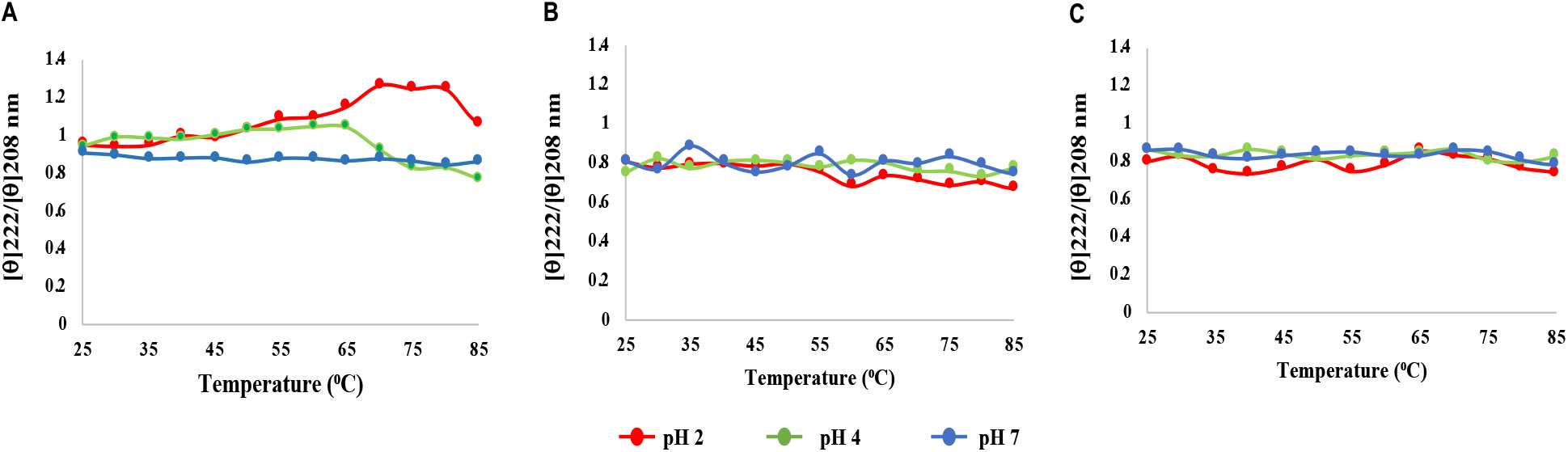
Ratio plots for HSP subunit CD data. Plots of [θ]222/[θ]208 nm as a function of temperature and pH for (A) HSPα, (B) HSPβ, and (C) HSPβ-coh.

### Fluctuations in pH and temperature induce changes in the tertiary structure of HSP subunits

Intrinsic protein fluorescence, predominately due to the fluorescent emission of tryptophan when excited at 280 nm, is a useful method to probe tertiary structural changes in proteins by providing information on stability and folding/unfolding states (47,48). Tryptophan fluorescence is sensitive to solvent polarity. Tryptophan residue(s) present on the surface of the protein are exposed to polar solvent whereas tryptophan residue(s) buried deep within a protein are shielded away from the polar environment. As the polarity of solvent increases, typically the fluorescence maxima goes through a red shift and the quantum yield decreases (49). However, in some cases, the tryptophan fluorescence of a protein may be quenched due to quenching by other residues (49). HSPα does not contain any tryptophan residues. The native conformation shows an emission maximum at 306 nm corresponding to the seven tyrosine residues in the protein located across all three domains: Y280, Y351 (apical); Y198, Y496 (intermediate); and Y124, Y128, Y430 (equatorial). For intrinsic fluorescence readings, HSPα was subjected to a pH range of 2-9 and held at temperatures 75°C, 80°C, and 90°C (Fig. 7A). Intrinsic fluorescence spectra of HSPα, across the pH range, show a maximum emission at ~ 306 nm, which is consistent with the native conformation of the protein (Fig. 7A). The environment of the fluorophores for HSPα remained unchanged as the emission maxima fluctuated only slightly by 2-4 nm with the exception of HSPα at 75°C and the higher pH range (pH 8 and 9) wherein the protein shows emission maxima shift closer to 350 nm. However, the fluorescence spectra of HSPα, under most conditions, show a small hump at 350 nm in addition to the more prominent emission maxima peak at 306 nm. This suggests that a subtle perturbation of the tertiary structure occurs due to temperature and pH fluctuations. The relative fluorescence intensity (RFI) is lower at lower pH conditions (pH 2 and 3) across all temperatures (physiological and heat shock). The decrease in intensity could be associated with an increase in compactness (supercoiling of helices), therefore resulting in a more buried environment of the fluorophores. This would decrease the exposure to solvent therefore decreasing emission. Overall, temperature and pH conditions did not lead to significant perturbations in the tertiary structure of HSPα.

**Figure 7.**
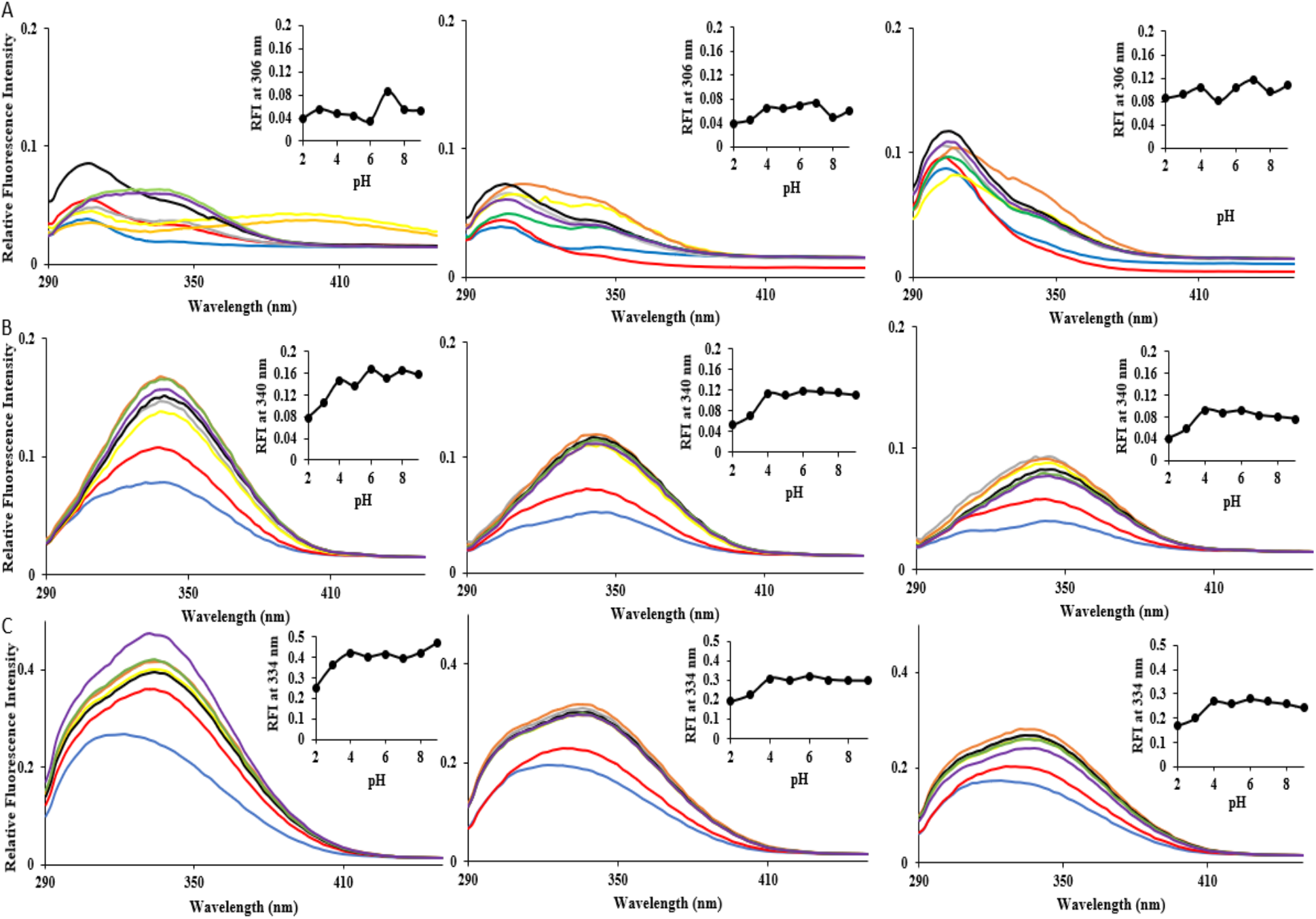
Intrinsic Fluorescence spectra of HSP subunits. Intrinsic Fluorescence (IF) spectra of each HSP subunit: (A) HSPα; (B), HSPβ, and (C) HSPβ-coh - subjected to a pH range of 2-9 at 75°C (left column), 80°C (middle column), and 90°C (right column top and middle for HSPα and HSPβ, respectively), and 85°C (right column bottom for HSPβ-coh) (right column). Inset graphs depict the changes in relative fluorescence intensity at the respective native emission maxima (nm) for each subunit as a function of pH. Color coding is the same for each graph: pH 2 (blue), pH 3 (red), pH 4 (grey), pH 5 (yellow), pH 6 (orange), pH 7 (black), pH 8 (green), pH 9 (purple).

HSPβ contains three tryptophan residues and 12 tyrosine residues located across the three domains: Y310, Y352 (apical); W198, Y199, Y223 (intermediate); and Y26, Y49, Y122, Y134, Y431, W433, Y446, Y484, W497 (equatorial). The emission maximum is at 340 nm in neutral pH over the range of temperatures (75°C, 80°C, and 90°C) suggesting that the folded state of the protein is relatively stable and that tryptophan residues for HSPβ are partially accessible to solvent in its native conformation. No significant changes in emission maxima were observed for HSPβ over this range of pH at 75°C (Fig. 7B) except for at pH 3 wherein the emission maximum decreased to 332 nm. At 80°C, the emission maximum for all pH conditions except for neutral pH (pH 7) shifted slightly by 2 nm. At 90°C, all emission maximum values went through a slight red shift of 2-4 nm. Again, the observed RFI is lower at pH 2 and 3 across all temperatures. As the emission maximum decreases, this suggests the fluorophores may have shifted to a more apolar core perhaps due to helical content increase.

Like HSPβ, HSPβ-coh also only contains three tryptophan residues. No additional tryptophan residues are introduced due to fusion of the cohesin to the HSPβ circular permutant. The emission maximum for HSPβ-coh is ~334 nm in neutral pH over the range of temperatures (75°C, 80°C, and 90°C). Interestingly, at pH 2 and 3, and over the full range temperatures (75°C, 80°C, and 90°C), HSPβ-coh exhibits a blue shift, as well as a lowered RFI consistent with the other HSPs (Fig. 7C).

Anilino naphthalene 8-sulfonate (ANS) is a non-polar dye which is used to probe the presence/extent of solvent-exposed hydrophobic surfaces in protein (41,50,51). This spectrophotometric technique is useful in the study of protein folding under different conditions. The degree of folding, the presence of potential intermediate states, or protein denaturation can be monitored by the exposure of hydrophobic residues on the surface of the protein. Differences in surface hydrophobicity of natural and engineered HSPs appear to be pH dependent (Fig. 8). ANS binding curves for HSPα and HSPβ at pH 7 and 75°C are comparable (i.e., superimpose). Furthermore, ANS binding for HSPα and HSPβ at pH 7 and 90°C (i.e., heat shock conditions) exhibit robust congruency. These results suggest that at both physiological pH and under heat shock, the native tertiary folds of these proteins are not significantly perturbed. ANS for HSPβ-coh at pH 7, under heat shock (i.e., 85°C) showed a slight increase in RFI as compared to the binding curve at physiological pH and temperature. For all HSPs at neutral pH and higher temperatures (75°C, 85°C, and 90°C), RFI for ANS binding is lower compared to the RFI at pH 2.

**Figure 8.**
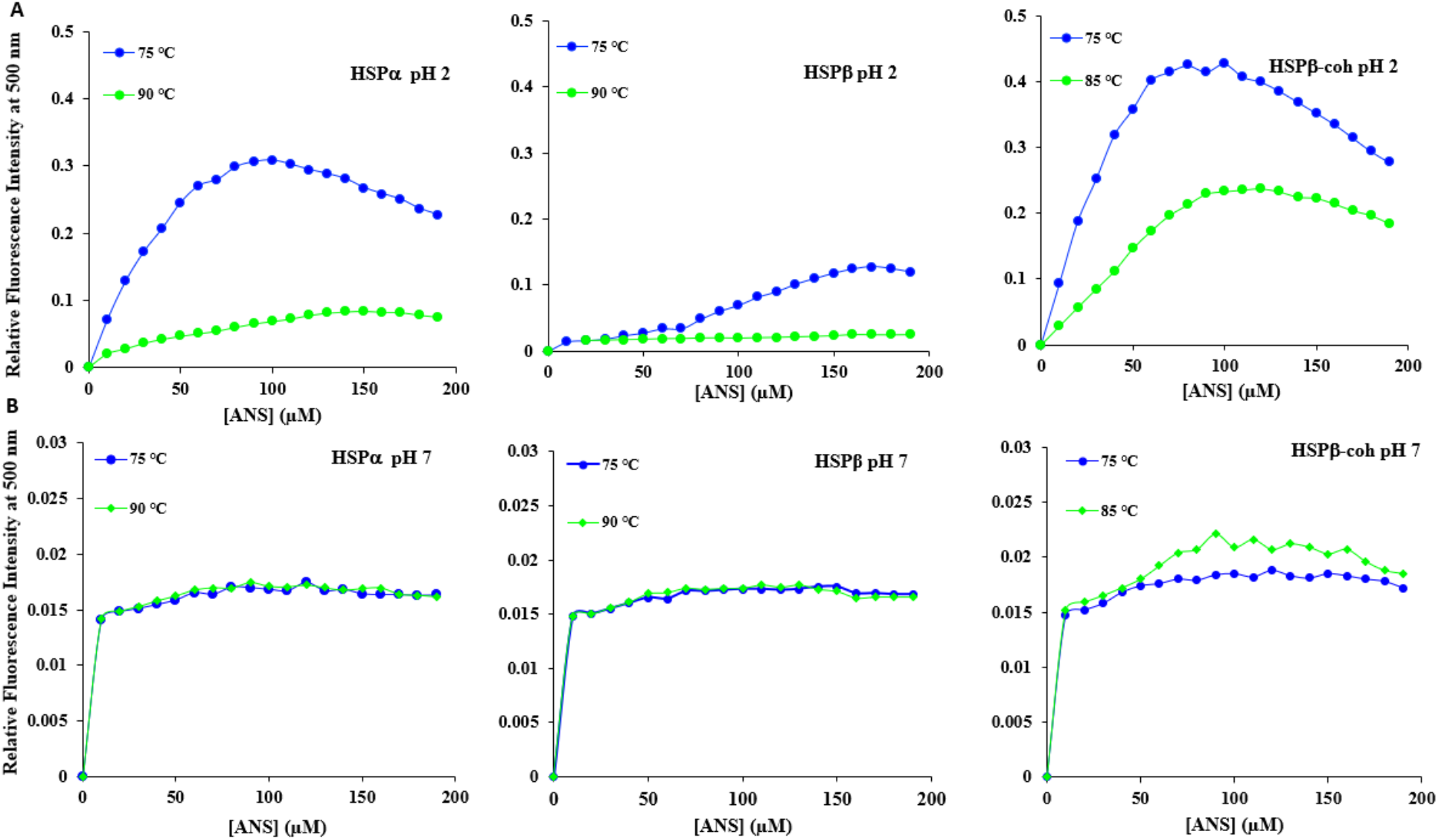
Anilino naphthalene 8-sulfonate (ANS) binding for each HSP subunit. ANS binding curves at: (A) pH 2 for HSPα (left), HSPβ (middle), and HSPβ-coh (right); and (B) pH 7 for HSPα (left), HSPβ (middle), and HSPβ-coh (right) - under two temperature conditions 75°C (blue) and 85° (HSPβ-coh, green) or 90°C (HSPα, HSPβ, green).

The hydropathic indices for each protein were calculated by the method of Kite and Doolittle (1982) and the values are averaged at each equivalent position (Fig. S3). Characterization of hydropathy profiles for natural and engineered HSP subunits regions revealed that the natural HSPα and HSPβ subunits were close mirror images of one another. The mean of the grand average hydropathy score (GRAVY) for natural HSPα, HSPβ, and the engineered fusion construct HSPβ-coh were calculated. These GRAVY scores are: 0.00625 for HSPα; −0.2 for HSPβ; and −0.11 for HSPβ-coh, respectively. Although, the hydrophilic profile of these proteins appears conserved across key structural elements, the hydrophobic properties around positions 550-700 of HSPβ-coh was shifted downward (i.e., less hydrophobic) when compared to the natural HSPαand HSPβsubunits.

### Limited trypsin digestion suggests a flexible HSPβ backbone structure

Trypsin, a serine protease, which cleaves proteins at the C-terminal end of lysine and arginine residues, is commonly used to assess the backbone flexibility of proteins and to provide low-resolution information regarding structural changes. Time-dependent trypsin digestion was employed to compare the backbone flexibilities between HSP subunits. The percent digestion post-incubation with trypsin for each HSP was measured by densitometric analysis of the 60 and 73 kDa bands in SDS PAGE gels (Fig.S4). The HSPβ-coh subunit is readily degraded by trypsin. Specifically, 87% of the protein is digested within 2 minutes of trypsin treatment. After 15 minutes of incubation with trypsin (37°C), the 73 kDa band corresponding to HSPβ-coh fusion protein is completely digested. In comparison, HSPβ and HSPα are digested to 75% and 60%, respectively, after 4 minutes (Fig. 9).

**Figure 9.**
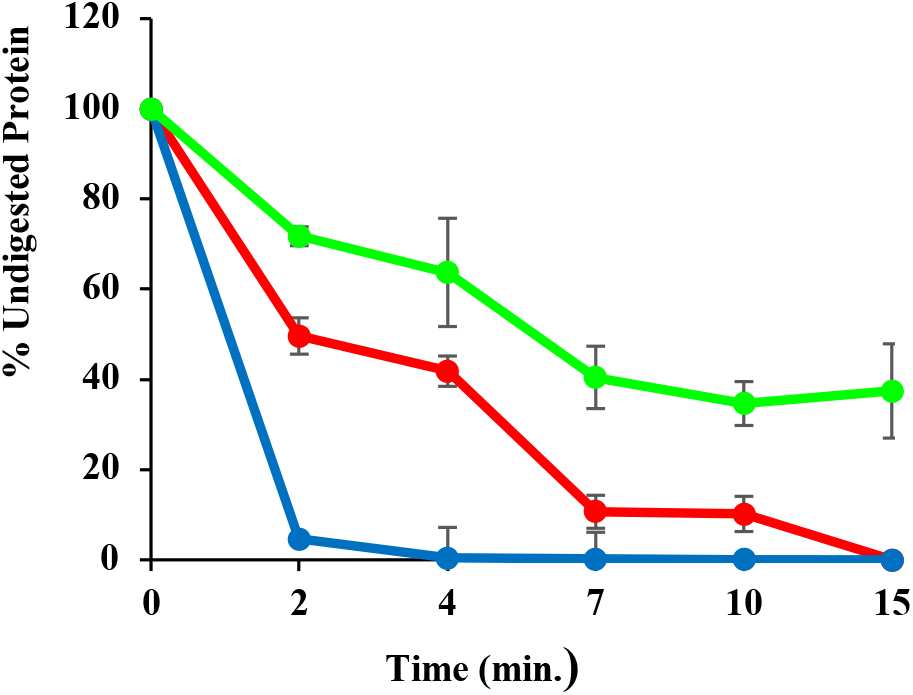
Densitometry for SDS-PAGE gels. depicting the resistance to limited trypsin digestion for HSPα (red), HSPβ (green), and HSPβ-Coh (Blue).

## Discussion

The structure (and function) of archaeal group II chaperonins has been previously studied (21,22,52). However, neither the staging of 18-mer HSP complex formation, the nature of HSP-client interactions, nor the extent of HSP complex protective action under extremes of pH and temperature is resolved. Unlike GroEL/ES, which uses a barrel (GroEL)-cap (GroES) system to serve clients, it is proposed that the group II chaperonin complexes (i.e., thermosomes) have both opened and closed states (18). A conformational change in the HSP complex allows client protein interactions within an inner cavity of the thermosome. It is the interaction of the client protein and the interior of the HSP complex that stabilizes the former under conditions of cellular stress (53–56). However, the conformational changes occurring at the level of the HSP subunit driving conformational changes in the complex are not clear. Likewise, the conformational states inducing subunits to form chaperonin complexes remain elusive. Although the upregulation of HSP subunit expression at physiological pH has been previously shown under heat shock conditions (55,57,58), recent evidence of HSP complex function in protecting client proteins under acidic (30,59) and alkaline (30) conditions suggests that HSPs may be resilient to changes in pH. Neither the extent to which natural or engineered HSPs maintain structural integrity at non-physiological pH nor the impacts of pH by temperature on subunit structure has been previously reported. Biophysical characterization of the limits to which HSPs can withstand changes in pH and temperature is necessary for designing engineered HSP platforms (i.e., MESP) for industrial applications as well as understanding the formation and function of natural HSP complexes (30).

### Natural and Engineered HSPs show thermostability

A gross measure of thermal stability can be determined by differential scanning calorimetry (DSC). The DSC output is a melting temperature (T_m_) at which 50% of the protein population is thermally denatured (40). All three HSP subunits exhibited significant thermostability. This aligns with previous work demonstrating HSP stability under conditions of thermal shock (18,21). HSPα and HSPβ were the most stable with T_m_ values at ~94°C at neutral pH (pH 7). The engineered HSPβ-coh subunit features a lower T_m_ of ~88°C at pH 7, which is still high compared to mesophilic proteins (60), and yet significantly lower than the T_m_ for the natural HSPα and HSPβsubunits. Given the similarities between the natural HSPβ chaperonins and the HSPβ circular permutant of the HSPβ-coh fusion construct, the lower T_m_ is likely due to thermal sensitivity in the fused cohesin domain. Although *C. thermocellum* is thermophilic (61,62), it is not a *hyper*thermophile; therefore, cohesin has not evolved at the same temperatures as group II chaperonins of the Sulfolobales (i.e., HSPα and HSPβ). There is a striking drop in T_m_ with decrease in pH. For all three HSP subunits, T_m_ drops ~25°C-30°C. This indicates a strong pH by temperature impact on the thermostability of these chaperonin subunits. To better understand the nature of this sensitivity to pH intrinsic fluorescence (IF), trypsin digestion, Anilino naphthalene 8-sulfonate (ANS), and circular dichroism (CD) were employed.

### Integrity of HSP tertiary structure shows pH dependency

To further explore pH by temperature effects on HSP structural stability, intrinsic tryptophan (Trp) and tyrosine (Tyr) fluorescence was used. Since the HSPα subunit does not contain Trp residues, IF for HSPα relied on Tyr-based fluorescence. IF_Tyr_ for HSPα shows a peak at 306 nm with the highest relative fluorescence intensity (RFI) at pH 7 under all temperature conditions. Although the IF_Tyr_ emission peak does not significantly shift upon altering pH, lower pH conditions result in decreased RFI compared to pH 7 across all temperature conditions indicating the possibility of supercoiling which was substantiated by the 222/208 nm ratio results for HSPα at pH 2. What is clear is that pH does impact the native tertiary fold. In HSPβ, IF_Trp_ shows an emission maximum at 340 nm, indicating partial exposure of tryptophan residues.

Multiple trials (including data from Fig. 7) suggest that low pH environments do not result in notable shifts in the emission maxima for HSPα and HSPβ across all temperature conditions. However, under all temperature conditions, decreases in RFI are observed most prominently at lower pH conditions (pH 2 and 3). HSPβ-coh exhibits a broader IF spectrum, possibly due to the additional tyrosine residues in cohesin that augments those of HSPβ. Likewise, since the HSPβ-coh is comprised of a circular permutant of HSPβ, the location of tryptophan residues is altered. Together, this likely account for the peak at ~334 nm and the less prominent shoulder at ~306 nm. A notable blue-shift is observed at pH 2 and 3 for HSPβ-coh at all temperatures examined, suggesting that ultra-low pH conditions cause this fusion protein to undergo changes in tertiary structure, which may indicate an unfolded intermediate state (Fig. 7, panel C).

DSC indicates a global breakdown in thermostability of HSPs under low pH conditions while IF indicates more subtle changes in HSP subunit conformation in pH by temperature interactions. To further investigate the effects of pH (and temperature) on tertiary structure, Anilino naphthalene 8-sulfonate (ANS) and trypsin digestion assays were conducted. ANS binding assays clearly demonstrate pH by temperature perturbations in HSP tertiary structure. At pH 7, ANS shows negligible changes in emissions from non-polar surfaces of HSPα and HSPβ at 75°C (physiological) and 90°C (heat shock). This result reflects that no significant changes to the solvent exposed hydrophobic surface(s) have been induced as a consequence of heat shock. At pH 7, HSPβ-coh shows minor increases in RFI with mild heat shock (85°C) the 75°C condition. (Again, the increase in RFI may be a result of ANS binding to the cohesion domain of HSPβ-coh). However, when the environment is adjusted to pH 2, perturbations in tertiary structure are observed. Specifically, significant increases in ANS RFI (i.e., 10-fold^+^) are observed for all three HSP subunits. This suggests that low pH conditions induce conformational changes in these chaperonins exposing non-polar surfaces to ANS binding. Since hydropathy plots indicate that HSPβ-coh features more hydrophobic regions than HSPα or HSPβ, increased ANS RFI amplitudes for HSPβ-coh is reasonable. Limited trypsin digest (LTD) is another method for assessing protein backbone stability/flexibility. Acknowledging that HSPβ-coh has more trypsin cleavage sites than HSPα or HSPβ: 82, 72, and 75, respectively (63). HSPβ has the highest backbone stability and therefore the least flexible structure of the three subunits. HSPα was slightly more susceptible to cleavage than HSPβ.The backbone structure of HSPβ-coh was observed to have the highest flexibility with the most susceptibility to cleavage, likely due to the addition of the cohesion moiety (see Fig. 7). Although increased thermostability and increased flexibility can coincide, thermophilic proteins have been shown to retain less flexibility than comparable mesophiles (64,65). Although LTD can only be done at neutral pH (to avoid disrupting enzyme activity), these data support results from IF, DSC, and ANS, suggesting that HSPβ is overall the most stable subunit under the pH and temperature conditions tested. This may be due to a more compact three-dimensional structure.

### HSP secondary structure exhibits resilience to shifts in pH

Although DSC, IF, and ANS data suggest significant pH by temperature effects on the structure of HSPs, these methods provide data on gross protein structural changes. It is critical to know whether the conformational shifts observed using these biophysical and biochemical techniques predominantly represent perturbations to tertiary structure or whether secondary structural elements are also impacted by pH. To explore the resilience of HSP secondary structure to changes in pH and temperature, circular dichroism (CD) was employed. All three HSPs tested show resiliency in the integrity of secondary structural elements over a range of temperatures (25°C-85°C) as demonstrated by far-UV CD. Taken together, CD and DSC data suggest that at physiological pH, which represents the pH of the intracellular environment of the Sulfolobus cell from which these subunits were derived, thermal stability is robust.

Comparisons of far-UV spectral data using both [θ]222/[θ]208 ratio and weighted spectral differences (WSD) indicate that secondary structural elements are stable across temperature ranges at neutral pH. Secondary structure also appears to be more resilient at pH 4 than pH 2 for all three subunits, with HSPα, exhibiting the greatest sensitivity to pH. The apparent stability of secondary structural elements at low pH with global collapse at ultra-low pH (e.g., pH 2) suggests that HSP tertiary structure is perturbed while preserving secondary structure under mildly acid (pH 3-6) conditions. Acid tolerance of secondary structure would allow for protein re-folding and preservation of function, while denaturation would likely occur at ultra-low pH. Whether the conformational shifts shown under mildly acid conditions represent intermediate conformational states associated with chaperonin-client protein interactions or HSP complex formation is not known. However, the presence of coiled-coil signatures in CD data suggest adaptation of a more compact structure at low pH.

### HSP gel data and structural models indicate pH by temperature conformational shifts

Protein gel electrophoresis that does not include use of a denaturant (i.e., SDS) may be described as Native PAGE (66). Native PAGE for purified HSPs at physiological pH reveals several bands including a high molecular weight band at ~1200 kDa likely representing the 18-mer complex. Depending upon whether ATP and MgCl_2_ is added to solution bands form with varying densities at molecular weights consistent with dimer, trimer, and nanomer (i.e., ring) formation. When HSP samples are subjected to ultra-low pH (i.e., pH 2) prior to conducting Native PAGE, gels resolve bands in the trimer and tetramer range for the ~60 kDA subunits. Taken with the CD data, these data raise the possibility that HSP conformational transitions (i.e., formation of coiled coils) may be essential for HSP complex formation. Specifically, interactions between α-helices on separate HSP molecules may comprise a fraction of the coiled-coil population. Movement between structural domains (i.e., apical, intermediate, equatorial) was observed in molecular dynamics simulations of HSPαand HSPβat pH 2 and pH 7 (70). Whether these shifts in tertiary structure are conformational intermediates required for HSP complex formation or HSP-client protein interaction requires further investigation.

### Experimental Procedures

#### Expression and Purification of Natural and Engineered HSP proteins

HSPα,HSPβ, and engineered HSPβ–cohesin fusion proteins were purified as previously described methods with some modifications (28,36). *Escherichia coli* expressing natural (HSPα, HSPβ and engineered (HSPβ-coh) subunits from *S. shibatae* were harvested by centrifugation at 7000 rpm for 15 min at 4°C (Sorvall RC-5) and cells were resuspended in buffer (50mM Tris-HCl, 1mM EDTA and pH 7.5), sonicated for 25 cycles (10 sec on/10 sec off) using SONIFIER 250, duty cycle 20%, and centrifuged at 16,000 x*g* for 20 min to separate supernatant from cellular debris. The supernatant extract was heated to 80 °C (HSPα and HSPβ) or 70 °C (engineered HSPβ-coh) for 30 min and centrifuged at 18,500 x*g* for 30 min to remove non-heat tolerant proteins. The supernatants were further purified to homogeneity by anion exchange chromatography (Bio-Rad Q Sepharose Column, USA), using a 150 mM to 1 M NaCl gradient for 30 min at 3 mL/min. The eluted proteins were collected and concentrated using a 30 kDa Amicon Ultra-15 centrifugal filter (EMD Millipore) and dialyzed overnight with 10 kDa SnakeSkinTM dialysis tubing (Thermo Scientific) at 4°C in 20 mM Tris–HCl (pH 8.0). All purified proteins were stored at −80°C for further analysis. The eluted fractions of protein concentrations were determined by Bradford protein estimation with BSA standards and proteins were analyzed by using SDS PAGE.

### Gel Electrophoresis (SDS-PAGE)

Proteins were resolved using 12% acrylamide resolving gel (Sambrook et al., 1989). Protein samples were mixed with SDS sample buffer (0.25% Coomassie Brilliant blue (R250), 2% SDS, 10 % glycerol (v/v), 100 mM tris, and 1% β-mercaptoethanol) at a ratio of 4:1 and boiled for 5 minutes. The boiled samples were loaded in respective wells and prestained protein molecular weight markers were also loaded (ThermoFisher Scientific, Cat.26620). The electrophoresis was performed at 150 volts for one hour using the Bio-Rad Mini protein electrophoresis system (Biorad, U.S.A). Gels were stained with Coomassie Brilliant Blue R-250 and destained with a mixture of methanol, water, and acetic acid (4:4:1).

### Silver staining of polyacrylamide gels

After electrophoresis, gels were incubated for 10 minutes in fixing solution (40% (v/v) methanol, 13.5% (v/v) formalin), resulting in precipitation of the proteins and diffusion of SDS. Subsequently, gels were placed into incubation solution (0.01g Na_2_S_2_O_3_) for 1 minute to oxidize the proteins. Gels were then washed with water three times for 5 minutes and transferred into silver solution (0.05 g silver nitrate) for another 10 minutes. Thereafter, proteins were visualized by replacement of the silver solution with developing solution (1.5 g Na_2_CO_3_, 25 μl formalin, 50 μl of 0.02% Na_2_S_2_O_3_). The sodium carbonate of the latter solution reduces the silver nitrate attached to the proteins and thus the proteins adopt a brown color. As soon as the desired staining intensity was reached, the reaction was stopped by addition of citric acid solution (2.3M), which was exchanged with water after 10 minutes.

### Circular dichroism spectroscopy

Far-UV circular dichroism (Far-UV CD) experiments were performed on a Jasco-1500 spectrophotometer. Thermal denaturation experiments were performed at temperatures ranging from 25-90°C at 5°C increments using protein concentrations of 0.3 mg/mL in 10 mM sodium phosphate buffer containing 10 mM NaCl with pH adjusted to 2, 4 and 7. The data was smoothened via application of the Savitzky-Golay algorithm.

The Weighted Spectral Difference (WSD) for each temperature and pH condition was calculated according to Dinh *et al.*, 2014 (43). The equation used to find the magnitude-weighted Euclidean distance between spectra was:

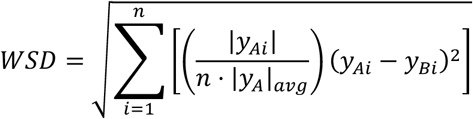

where *n* is the number of wavelengths measured within the spectral range, *y*_*Ai*_ is the *i*^th^ sample of the reference spectra, *y*_*Ai*_ is the ith sample of the sample spectra, and |*y*_*A*_|_*avg*_ is the average absolute molar ellipticity of the reference sample. The smoothed spectra corresponding to pH = 7 and T = 75°C was used as the reference spectra for each protein. The range of wavelengths 200 to 250 nm were utilized in this calculation. All WSD calculations were completed in R (R_CS).

### Fluorescence spectroscopy and 8-Anilino-1-napthalenesulfonic acid (ANS) binding

All intrinsic fluorescence measurements were performed on a Jasco-1500 spectrophotometer using a 10 mm quartz cuvette with protein concentrations of 0.2 mg/mL in 10 mM sodium phosphate buffer containing 100 mM NaCl with pH ranging from 2-9 at pH increments of 1 and held at temperatures 75°C, 80°C, 85°C, and 90°C. Protein samples were excited at a wavelength of 280 nm and the emission spectra was recorded from 290 nm to 450 nm. ANS binding assays were performed using protein concentrations of 0.2 mg/ml. Stock ANS was made such that 2 μL titrations into the protein sample would increase the ANS concentration by 10 μM increments. Fluorescence measurements were made after each titration until saturation was reached. Samples were excited at 380 nm and emission was recorded at 500 nm. Measurements were recorded under variable pH buffer conditions (pH 2 and 7) and variable temperature conditions (75°C, 85°C, and 90°C)

### Differential Scanning Calorimetry (DSC)

All protein samples were prepared at a concentration of 0.5 mg/ml in 10 mM phosphate buffer. A N-DSC III Differential Scanning calorimeter was used to determine the melting temperatures. Prior to loading, all the protein samples were subjected to degassing at 25°C for 15 minutes. Scans were performed from 25 to 110 °C with a 1 °C/min ramping temperature and at variable pH solutions (pH 2, 4, and 7). To obtain a stable baseline, buffer runs were conducted before running the protein scans. Blank subtraction was performed, and data obtained was processed using CpCalc Version 2.2.0.10 software provided by the manufacturer.

### Limited Trypsin Digestion

Limited trypsin digestion experiments were performed on all HSP subunits. The initial reaction mixture included 0.2 mg/ml of protein and 0.000781 mg/ml of trypsin in 10 mM sodium phosphate buffer at pH 7.2. Trypsin digestion was carried out at 37 °C and a portion of the reaction mixture was removed, at specified time intervals over 15 minutes. The reaction was arrested by precipitation using trichloroacetic acid. The reaction products were analyzed by 15% SDS-PAGE and the gels were stained using Coomassie Brilliant Blue (Sigma Aldrich). The band intensities for SDS-PAGE were estimated to determine the percentage of proteolytic digestion using UN-SCAN-IT densiometric software. The intensity of HSP samples not subjected to proteolytic digestion was used as the control representing 100% protection from enzymatic degradation.

### Structural Modeling

The FASTA sequences of the Heat Shock Proteins (HSPα and HSPβ) from *S.shibatae*, were uploaded to the SWISS Model (67). Sequences with similarity of 60% or more were chosen for the templates of HSPα and HSPβ, to generate 3-dimensional models in PDB format for both proteins. After that, the chosen models were subjected to energy minimization and RMSD Calculations using Swiss PDB Deep Viewer to select the best model for each protein (68). Further, HSPα and HSPβ were subjected to scan using EMBL-EBI InterPro to visualize the domains (34). The highly conserved domains among all the sub-units, including HSPβ-coh, were marked using ESPript (69). The Secondary Structures for the HSPs were visualized using the Swiss PDB Deep Viewer (68). NsitePred webserver was used to determine the possible ADP/ATP exchange sites (37). Finally, PyMOL (The PyMOL Molecular Graphics System, Version 2.0 Schrödinger, LLC.) was used to visualize and label the three functional domains (Apical, Intermediate, and Equatorial), the conserved domains, Secondary Structure (Alpha Helices, Beta Sheets and Turns), the hydrophobic residues, ATP sites, and the N or C Terminus in the 3-Dimensional structure of the proteins.

## Conclusions

Detailed biophysical analysis of natural chaperonins (HSPα and HSPβ) and an engineered HSPβ-coh construct indicate that, despite the high thermostability of all three subunits, the tertiary structures of all three HSPs are affected by pH. Other work by our lab has indicated that HSP complexes may be resilient to low pH conditions (71); however, biophysical analyses indicate that the tertiary structures of these HSPs are sensitive to changes in pH. Nonetheless, secondary structural elements of these HSPs do show some resilience to acid conditions suggesting that function may be retained after pH challenge. A molecular dynamics simulation study (70) supports this suggestion. Additional studies are required to demonstrate retention of HSP function after pH challenge. This is the subject of ongoing work in the lab. More work is required to explore conformational shifts in these HSP subunits to determine if intermediate states induced by pH by temperature challenges are important for HSP complex formation, HSP-client protein interactions, or switching between opened and closed states in a functional thermosome (i.e., 18-mer HSP complex). Detailed analyses of structural changes including potential intermediates to key conformational changes are required for both natural and engineered constructs to direct development of artificial HSP platforms (i.e., MESPs).

## Acknowledgements

This work was supported by two awards from the U.S. National Science Foundation: a NSF MCB grant (award no. 1818346; PI-Ceballos) and a NSF INFEWS grant (award no. 1856091; PI-Ceballos). A University of Arkansas Chancellor’s Innovation and Commercialization award supported the participation of MF and SB. The authors would like to thank Dr. Mary Cloninger of Montana State University and Dr. J.B. Alexander “Sandy” Ross of the University of Montana for comments and suggestions during the preparation of this manuscript.

