## Supplemental Data for "Stability Comparisons between Natural versus Engineered Archaeal Heat-Shock Proteins"

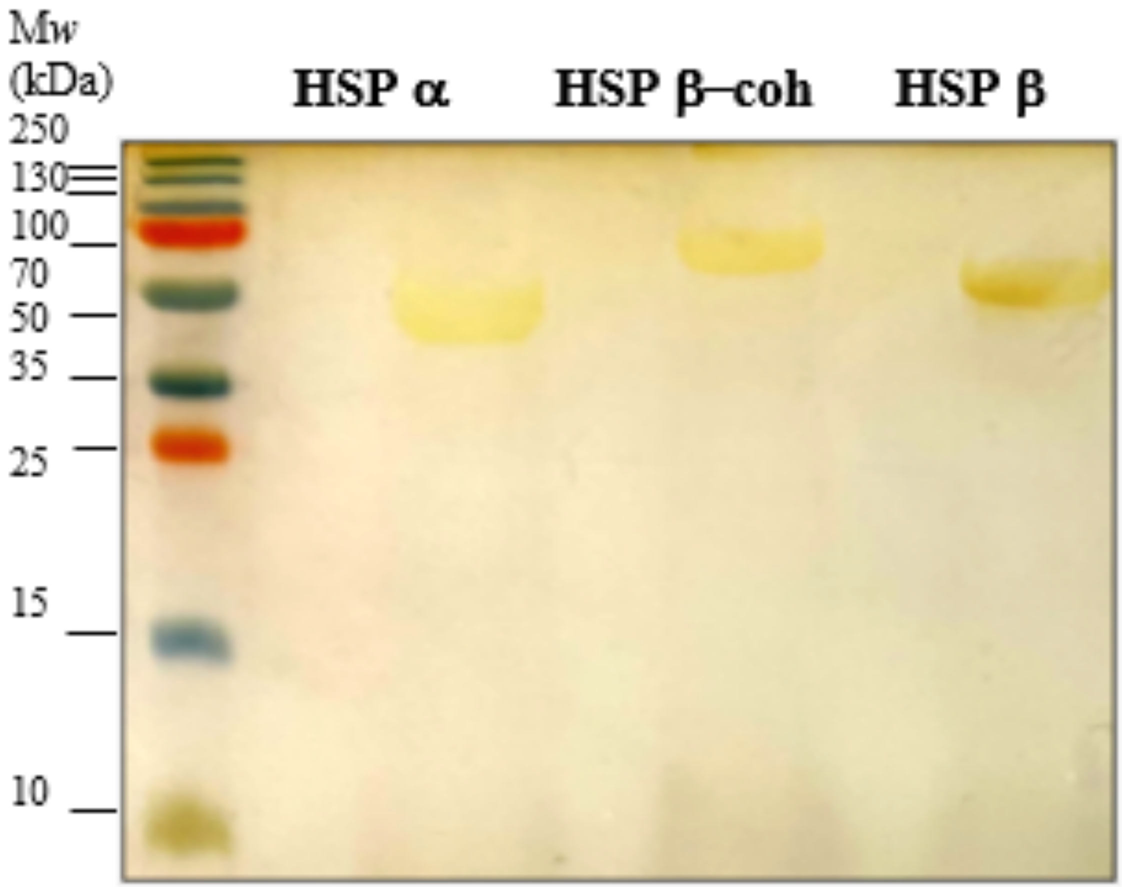


**Figure S1**. Purification of HSP natural (α and β) and engineered (β-coh) from the E. coli Codonplus/pET19b. The purity of HSP subunits is visualized on silver-stained SDS-PAGE

**HSPα**

MASPVLLLKEGTSRTTGRDALRNNILAAKTLAEMLRSSLGPKGLDKMLIDSFGDVTITNDGATIVKDMEIQHPAAKLLVEAAKAQDAEVGDGTTSAVVLAGALLEKAESLLDQ
NIHPTIIIEGYKKAYTKALELLPQLGTRIDIRDLNSSVARDTLRKIAFTTLASKFIAEGAELNKIIDMVIDAIVNVAEPLPNGGYNVSLDLIKIDKKKGGSIEDSVLVKGLVLDKEVVHP
GMPRRVTKAKIAVLDAALEVEKPEISAKISITSPEQIKAFLDEESKYLKDMVDKLASIGANVVICQKGIDDIAQHFLAKKGILAVRRVKRSDIEKLEKALGARIISSIKDATPDDLGYA
ELVEERRVGNDKMVFIEGAKNLKAVNILLRGSNDMALDEAERSINDALHALRNILLEPVILPGGGAIELELAMKLREYARSVGGKEQLAIEAFADALEEIPTILAETAGLEAISAL
MDLRARHAKGLTNTGVDVIGGKIVDDVYALNIIEPIRVKAQVLKSATEAATAILKIDDLIAAAPLKSEKKGGEGSKEESGGEGGAGTPSLGD

**HSPβ**

MATATVATTPEGIPVIILKEGSSRTYGKEALRANIAAVKAIEEALKSTYGPRGMDKMFVDSLGDITITNDGATILDKMDLQHPTGKLLVQIAKGQDEETADGTKTAVILAGELAKK
AEDLLYKEIHPTIIVSGYKKAEEIALKTIQDIAQPVSINDTDVLRKVALTSLGSKAVAGAREYLADLVVKAVAQVAELRGDKWYVDLDNVQIVKKHGGSINDTQLVYGIVVDKEVV
HPGMPKRIENAKIALLDASLEVEKPELDAEIRINDPTQMHKFLEEEENILKEKVDKIAATGANVVICQKGIDEVAQHYLAKKGILAVRRAKKSDLEKLARATGGRVISNIDELTSQ
DLGYAALVEERKVGEDKMVFVEGAKNPKSVSILIRGGLERVVDETERALRDALGTVADVIRDGRAVAGGGAVEIEIAKRLRKYAPQVGGKEQLAIEAYANAIEGLIMILAENAGL
DPIDKLMQLRSLHENETNKWYGLNLFTGNPEDMWKLGVIEPALVKMNAIKAATEAVTLVLRIDDIVAAGKKGGSEPGGKKEKEEKSSED

**HSPβ-coh**

MADPTQMHKFLEEEENILKEKVDKIAATGANVVIAQKGIDEVAQHYLAKKGILAVRRAKKSDLEKLARATGGRVISNIDELTSQDLGYAALVEERKVGEDKIVFVEGAKNPKSVS
ILIRGGLERVVDETERALRDALGTVADVIRDGRAVAGGGAVEIEIAKRLRKYAPQVGGKEQLAIEAYANAIEGLIMILAENAGLDPIDKLMQLRSLHENETNKWYGLNLFTGNPE
DMWKLGVIEPALVKMNAIKAATEAVTLVLRIDDIVGGSGGTIPVIILKEGSSRTYGKEALRANIAAVKAIEEALKSTYGPRGMDKILVDSLGDITITNDGATILDKMDLQHPTGKLL
VQIAKGQDEETADGTKTAVILAGELAKKAEDLLYKEIHPTIIVSGYKKAEEIALKTIQDIAQPVSINDTDVLRKVALTSLGSKAVAGAREYLADLVVKAVAQVAELRGDKWYVDLD
NVQIVKKHGGSINDTQLVYGIVVDKEVVHPGMPKRIENAKIALLDASLEVEKPELDAEIRINGGSGGSGGSVPSDGVVVEIGKVTGSVGTTVEIPVYFRGVPSKGIANCDFVFR
YDPNVLEIIGIDPGDIIVDPNPTKSFDTAIYPDRKIIVFLFAEDSGTGAYAITKDGVFAKIRATVKSSAPGYITFDEVGGFADNDLVEQKVSFIDGGVNVGNATP

**Figure S2.** Primary sequences of HSPα, HSPβ, and HSPβ-coh. Cohesin moiety is highlighted in green.

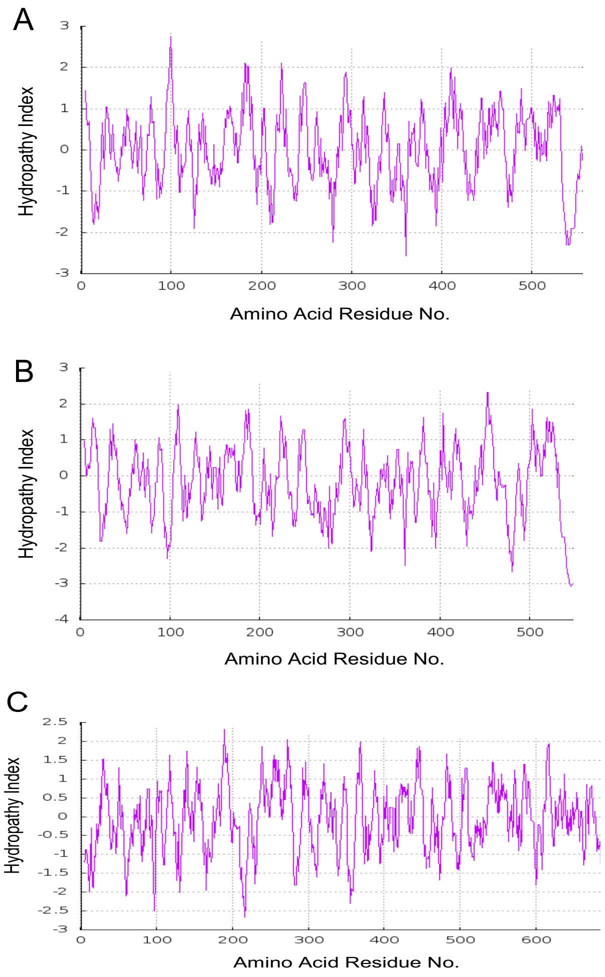


**Figure S3. Hydropathy Plots for HSP subunits.** Hydropathy indices of (A) HSPα, (B) HSPβ and (C) HSPβ-coh are generated by the by the ProtScale (Expasy) web server using the method of Kyte and Doolittle (1982), with positive (hydrophilic) and negative (hydrophobic) values plotted above and below the center lines, respectively.

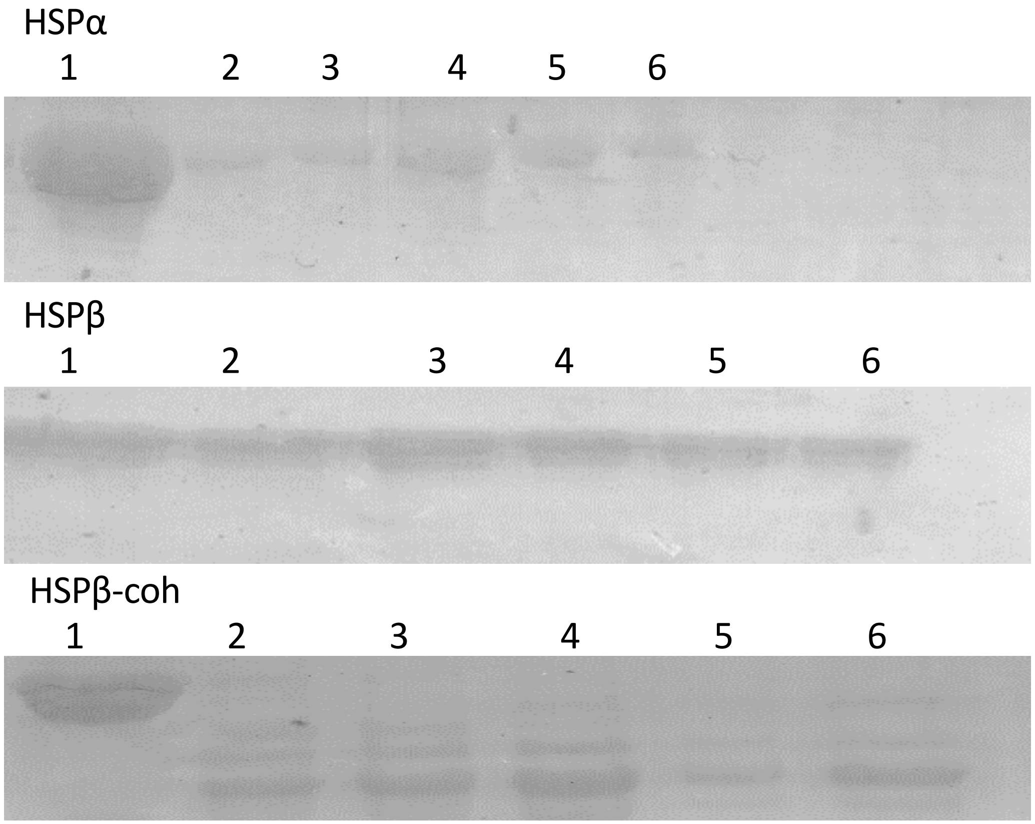


**Figure S4.** SDS PAGE analysis of the limited trypsin digestion of HSPα, HSPβ, and HSPβ-coh on SDS-PAGE. Each lane indicates the amount of undigested protein at the respective time intervals. Lane 1= 0 trypsin, Lane 2= 2 minutes incubation with trypsin, Lane 3= 4 minutes incubation with trypsin, Lane 4= 7 minutes incubation with trypsin, Lane 5= 10 minutes incubation with trypsin, Lane 7= 15 minutes incubation with trypsin.
